# iSeqsSearch: Incremental Protein Search for iBlast/iMMSeqs2/iDiamond

**DOI:** 10.1101/2024.09.09.612094

**Authors:** Hyunwoo Yoo, Mohammad S. Refahi, Robi Polikar, Bahrad A. Sokhansanj, James R. Brown, Gail L. Rosen

**Affiliations:** Department of Electrical and Computer Engineering, Drexel University, Pennsylvania, United States; Electrical and Computer Engineering, Rowan University, Engineering Hall 346B, 08028, New Jersey, United States

## Abstract

**Background:** The advancement of sequencing technology has led to a rapid increase in the amount of DNA and protein sequence data; consequently, the size of genomic and proteomic databases is constantly growing. As a result, database searches need to be continually updated to account for the new data being added. Continually re-searching the entire existing dataset, however, wastes resources. Incremental database search can address this problem.

**Methods:** One recently introduced incremental search method is iBlast, which wraps the BLAST sequence search method with an algorithm to reuse previously processed data and thereby increase search efficiency. The iBlast wrapper, however, must be generalized to support more performant DNA/protein sequence search methods that have been developed, namely MMseqs2 and Diamond. Moreover, the previously published iBlast wrapper has to be revised to be more robust and usable by the general community.

**Results:** iMMseqs2 and iDiamond, which apply the incremental approach, obtain results nearly identical to those achieved using only MMseqs2 and Diamond. Notably, when comparing ranking comparison methods such as the Pearson correlation, we observe a high concordance of over 0.9, indicating similar results. Moreover, in some cases, our incremental approach applying iBlast merge function and using m8 formats including the new m8e format provides more hits compared to the conventional MMseqs2 and Diamond.

**Conclusion:** The incremental approach using iMMseqs2 and iDiamond demonstrates efficiency in terms of reusing previously processed data while maintaining high accuracy and concordance in search results. This method can reduce resource waste in continually growing genomic and proteomic database searches. The sample codes are made available at GitHub: https://github.com/EESI/Incremental-Protein-Search.

## INTRODUCTION

In the realm of genomic and proteomic research, the advent of high-throughput sequencing technologies has precipitated an unprecedented expansion in the volume of nucleic acid and amino acid sequence data. This deluge of data has, in turn, necessitated the development and expansion of comprehensive sequence databases to catalog and make sense of this wealth of information. The traditional approach of exhaustive search within these ever-growing repositories, however, poses a significant challenge in terms of computational efficiency and resource allocation.

This rapid accumulation of sequence data is not merely a technical challenge but a fundamental shift in our ability to understand biological systems. The exponential growth of databases like UniProtKB/Swiss-Prot, which now contains millions of protein sequences, offers unprecedented opportunities for discovering evolutionary relationships, predicting protein structures, and understanding molecular functions. However, it also presents substantial computational hurdles that demand innovative solutions.

The incremental search method, which updates only new or modified data, was introduced through NBC++Zhao et al. (2020) which focus on incrementally processing the results of alignment search rather than presenting the results of clustering. Subsequently, an incremental method utilizing BLAST, known as iBlastDash et al. (2021), was also developed. They emerged as a solution to mitigate the computational burden by leveraging previously processed data within the framework of the BLAST Altschul et al. (1990) algorithm. Despite its utility, the advent of more advanced and efficient search tools such as Diamond Buchfink et al. (2021) and MMseqs2 Steinegger and Soöding (2017), which offer superior performance in handling large-scale sequence data, has rendered the traditional BLAST-based methods less optimal.

This paper introduces a methodology that integrates the incremental search principle of iBlast with the advanced search capabilities of Diamond and MMseqs2. (Note that MMSeqs2 has an incremental clustering ability **?**, but our paper focuses on incremental search). Our approach is designed to enhance search efficiency in the face of continuously expanding sequence databases, thereby supporting sustainable and effective data management and retrieval. Specifically, we present a novel approach that allows for rapid integration of new sequences into existing search results, without the need for complete database re-scans. Furthermore, we introduce an extended file format, m8e, which enhances the standard m8 format by incorporating critical metadata.

## MATERIALS & METHODS

We develop an incremental method for efficient protein sequence similarity searches, utilizing the Scope Astral protein database Fox et al. (2014) as our benchmark dataset. This study implements an incremental method that utilizes Spouge statistics Park et al. (2012) in iBlast, similar to the widely used iBlast method, to maintain result accuracy while improving the processing speed for new data. Our method employs a two-step process: (1) initial database search using BLAST, MMseqs2, or Diamond with default parameters, and (2) incremental updates using our custom algorithm based on Spouge statistics in iBlast. Instead of complex XML files, this process uses the m8 file format for data merging, simplifying the conversion and integration of these files. The m8 format contains tab-separated fields: query id, subject id, percent identity, alignment length, mismatches, gap opens, q.start, q.end, s.start, s.end, E-value, and bit score. Additionally, we propose an extended version of the m8 file format, herein named m8e (m8 extension). The m8e format includes the total length of the database within the file, adding an extra line it the beginning of each file to store this information. The database length is crucial for accurate E-value calculations during incremental updates. This allows for the automatic integration of two result files.

We implement our method in Python (version 3.8+), utilizing the Biopython library (version 1.79) for sequence handling and NumPy (version 1.21.0) for numerical operations. To evaluate the incremental method’s performance, we conduct comprehensive benchmarks comparing our incremental approach against full database searches. We divide the Scope Astral protein database into 10 batches, with the first batch stratified based on protein class and the remaining 9 batches all stratified in the same way as the first batch (and then the individual batches randomly partitioned). The first batch serves as the query set, while the remaining 9 batches were sequentially combined to form increasingly larger search databases, simulating database growth over time. Experiments use Blastp, MMseqs2, and DIAMOND with default settings unless otherwise stated (changing the hits returned under an E-value threshold and also extending the limit of the returned hits beyond the default limits), only adjusting the number of threads to 32 for consistency. For each database size (from 2 to 10 batches), we performed both full and incremental searches.

## RESULTS

Our analysis of the incremental method for protein sequence searches revealed improvements in both search effectiveness and computational efficiency compared to traditional non-incremental methods.

Fig. 1 a) illustrates two primary findings: First, incremental methods (iBlastp, iMMseqs2, iDiamond) consistently yield higher hit counts compared to their non-incremental counterparts (Blastp, MMseqs2, Diamond). This increase in hits suggests that incremental methods are capable of identifying additional potential matches, thereby enhancing the comprehensiveness of the search. However, not all methods are equal in their incremental search. Second, the processing times for incremental methods are substantially reduced, as shown in the right graph of Fig. 1 b). This time efficiency is crucial for large-scale protein database searches where computational resources are often a limiting factor.

**Figure 1.**
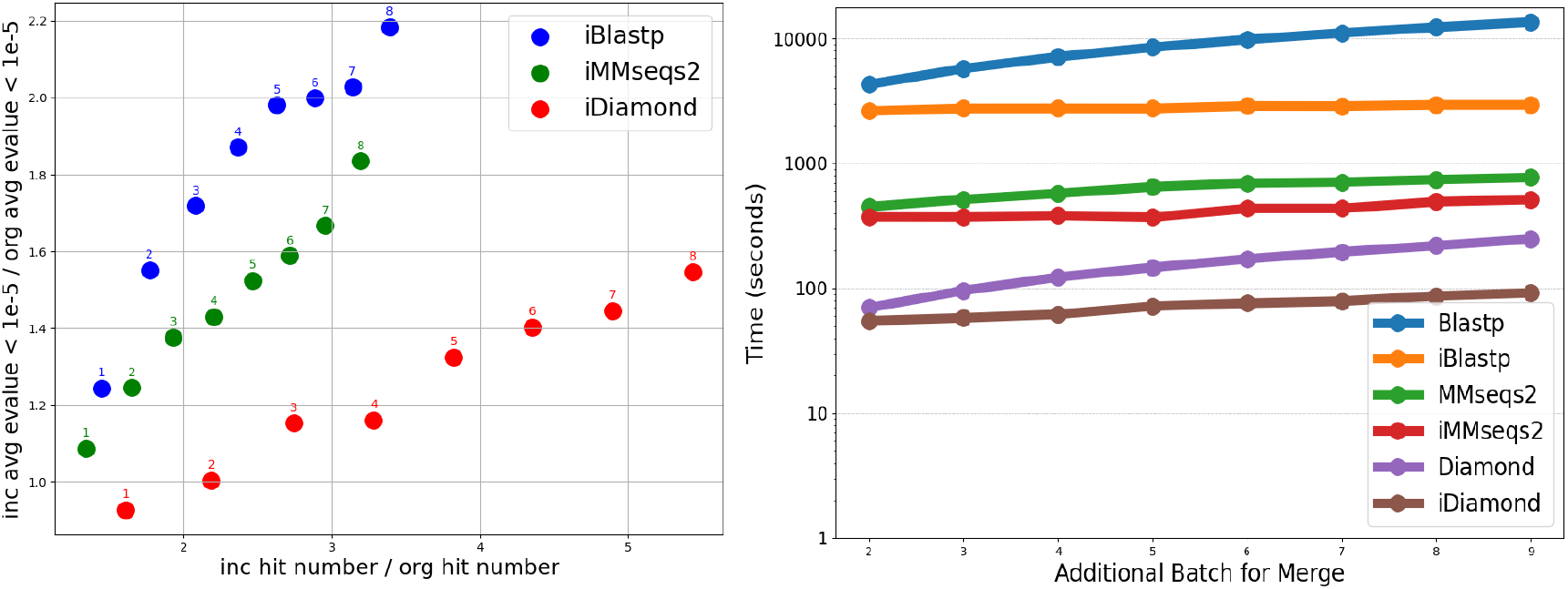
The left graph (a) illustrates increasing E-values and hit counts when comparing incremental methods (iBlastp, iMMseqs2, iDiamond) to non-incremental methods (Blastp, MMseqs2, Diamond). The right graph (b) compares the processing times of these methods, showing that incremental methods are faster.

It is important to note that while incremental methods result in increased hit counts, they also lead to larger E-values (Fig. 1 a). Although larger E-values typically indicate less statistically significant matches, we only include hits below the commonly used threshold of 1e-5 in the figure 1 a). This suggests that additional matches, while potentially less statistically confident, may still be biologically relevant and worthy of consideration in many applications. As a case study, we examine one type of protein for hits that have the “correct”/“right” protein family label vs. hits that have incorrect labels. In general, the incremental version had higher E-values than the non-incremental version. Compared to Blast and MMSeqs2 (Figs. 22 and 23), Diamond (Fig. 24) returns less hits on every batch and their E-value is much lower (more significant).

When incremental batch learning is used, there is a reduction in queries with no hits. This reduction further underscores the effectiveness of the incremental method. As shown in Figure 15 in the Supplementary Materials, the proportion of queries with zero hits using incremental methods is lower than that of non-incremental methods, indicating improved search coverage.

To assess the quality and consistency of search results, we show the results of several correlation measures. While there are several similarity measures available, including the Pearson correlation coefficient Pearson (1896), the Kendall tau correlation coefficient Kendall (1938), and the Spearman correlation coefficient Spearman (1904), this study specifically utilizes the 1) Pearson to assess the concordance of the search methods in their E-value and 2) the Kendall tau measure to assess the concordance between the methods’ rankings. Fig. 2 presents heatmaps of the Kendall tau and Pearson correlation coefficients for each search method. The Pearson correlation heatmap reveals high similarity between incremental and non-incremental methods in the E-values of overlapping hits between methods, with iBlastp and Blastp achieving a score of 0.97, while iMMseqs2 and MMseqs2, as well as iDiamond and Diamond, shows perfect correlation with a score of 1.0 (Fig. 2). These high correlation scores indicate that the incremental methods maintain result quality comparable to their non-incremental counterparts.

**Figure 2.**
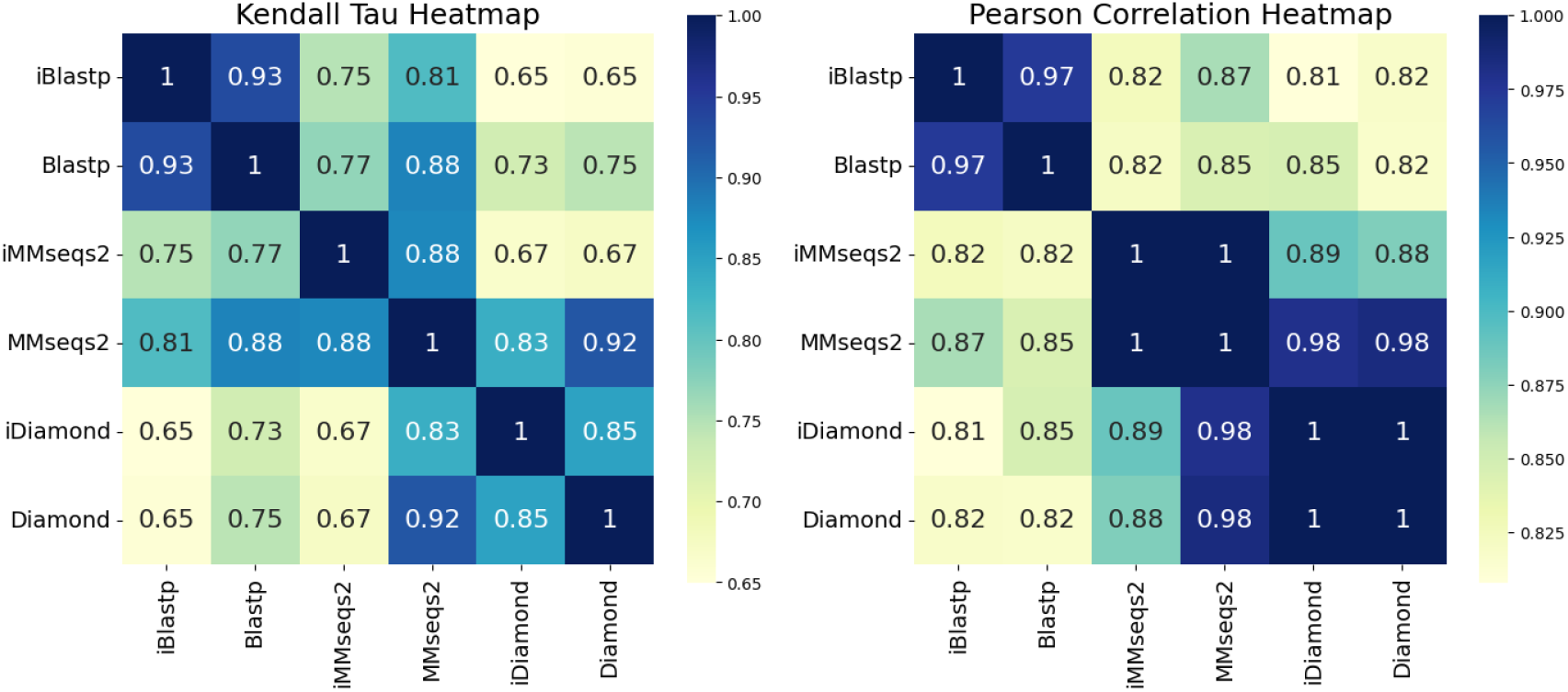
Heatmaps representing the Kendall tau correlation coefficient and the Pearson correlation coefficient for each search result. In the searches, 1/10th of the Astral Scope dataset is used as the query, and the remaining dataset is used as the database. The database is randomly sampled and divided into nine equal batches, which is then incrementally combined based on the search results of the fully combined dataset. In the Pearson correlation heatmap, iBlastp and Blastp achieves a score of 0.97, which was the highest among the methods, while iMMseqs2 and MMseqs2, as well as iDiamond and Diamond, shows a score of 1.0, indicating the highest similarity. For the Kendall tau correlation, iBlastp and Blastp scores 0.03, which was higher than the other methods, and iDiamond and Diamond also had a higher score of 0.85. This indicates that iBlastp and Blastp, as well as iMMseqs2 and MMseqs2, and iDiamond and Diamond, provide similar results.

The Kendall tau correlation, which measures rank consistency, also shows strong agreement between incremental and non-incremental methods. iBlastp and Blastp scores 0.57, while iDiamond and Diamond achieves 0.48, both higher than other method pairs (Fig. 2). These results suggest that the incremental methods preserve the ranking of hits, an important factor in sequence similarity searches.

The quality of the searches is also measured using log Discounted Cumulative Gain Jaärvelin and Kekaälaäinen (2002) and normalized Discounted Cumulative Gain Wang et al. (2013). We evaluate the search quality using log Discounted Cumulative Gain (DCG). The consistent increase in log DCG values across incremental methods indicates improvements in the quality of search results, reflecting both better ranking of hits and more effective identification of relevant protein sequences as shown in Supplementary Figure 16.

Furthermore, we compare the average number of hits per query for each method with and without maximum target hit limits (Table 1). iBlastp achieves a remarkably high average of 725.93 hits per query, surpassing both Blastp with no limit (253.58 hits) and with the default limit (240.89 hits). However, without a maximum limit, Diamond achieves an average of 435.78 hits per query higher than the 22.14 hits with the default limit and 120.38 hits with iDiamond (which use default Diamond parameters).

**Table 1.** Comparison of Average Hits(e-value is less than 1e-5) per Query and Time (seconds) for Different Methods. Average Hits per seconds indicates Average Hits per Query per seconds. Default limitations are 25, 300, 500 for Diamond, MMseqs2, Blastp.

| Method | Average Hits<br>per Query | Time<br>(seconds) | Average Hits<br>per Time |
| --- | --- | --- | --- |
| Blastp (No Limit) | 253.58 | 48000 | 0.0052 |
| Blastp (Default) | 240.89 | 13566 | 0.0178 |
| iBlastp (each batch avg.) | 446.05 | 2820 | 0.1581 |
| iBlastp (all batches) | 725.93 | 22566 | 0.0322 |
| MMseqs2 (No Limit) | 682.29 | 48160 | 0.0141 |
| MMseqs2 (Default) | 166.74 | 771 | 0.2163 |
| iMMseqs2 (each batch avg.) | 316.77 | 435 | 0.7282 |
| iMMseqs2 (all batches) | 517.08 | 3381 | 0.1529 |
| Diamond (No Limit) | 435.78 | 1792 | 0.2431 |
| Diamond (Default) | 22.14 | 249 | 0.0889 |
| iDiamond (each batch avg.) | 73.29 | 73 | 1.003 |
| iDiamond (all batches) | 120.38 | 581 | 0.2072 |

The efficiencies of incremental methods are further highlights by their reduced search times. For instance, iDiamond completes searches approximately 19 times faster than Diamond without a hit limit. iMMseqs2 was about 2 times faster, and iBlastp was around 16 times faster than their non-incremental counterparts. Venn diagram analysis (Figure 12, 13 and 14 in Supplementary Materials) reveals that over 90% of the hits from non-incremental methods are included in the hits from incremental methods, demonstrating that incremental methods maintain comprehensive coverage while improving efficiency.

We also aim to see how well we could classify the queries into their 7 SCOPe protein classes after each batch. The graphs (Figures 9, 10, and 11 in Supplementary Material) compare the protein class F1 score of non-incremental methods and incremental methods based on the E-value top hit criterion. The results of the incremental experiments show a trend of increasing F1 score across all cases, with classification linearly increasing as data is added up to 99% when all SCOPe classes are known. We also observe that the trend of increasing F1 scores was similar for both incremental and non-incremental methods. While the performance of incremental methods was sometimes slightly better (e.g., iMMseqs2) or slightly worse (e.g., iDiamond), the overall trend remains similar.

## DISCUSSION

This study confirmed that incremental methods in protein sequence searches provide broader coverage by generating more hits than non-incremental methods. Additionally, by using Discounted Cumulative Gain (DCG), a ranking metric often used in recommendation systems, we verified that the additional hits identified by incremental methods are biologically meaningful. This suggests that incremental methods not only increase the number of hits but also yield qualitatively significant results.

The reduced processing times of incremental methods offer substantial advantages, particularly when searching large-scale databases. Future research should analyze the performance differences across databases of varying sizes. However, in this study, the comparison of performance based on database size is left for future work.

While there is a tendency for E-values to increase, we found that applying a 1e-5 threshold provided effective filtering. This demonstrates that incremental methods, while yielding more hits, offer results that are comparable in reliability to those of non-incremental methods. Therefore, incremental methods can maintain statistically significant results while encompassing a broader range of hits.

In terms of protein SCOPe classification, incremental methods showed improved performance in some categories. Notably, iMMseqs2 achieved higher F1 scores, suggesting that incremental methods can also be useful for protein classification tasks. Future research should explore the applicability of incremental methods in other classification systems, such as CATH and Pfam.

## CONCLUSIONS

In conclusion, these results demonstrate that the incremental method implemented in Incremental-Protein-Search offers advantages over traditional sequence search methods. Incremental-Protein-Search increases hit counts, reduces computational time, and maintains result quality and ranking consistency. The trade-off between slightly higher E-values and increased hit counts should be carefully considered in the context of specific research goals. Higher E-values can lead to the inclusion of more false positives, which may lower the overall accuracy of the results. The incremental learning method applied to any base search enhances both efficiency and accuracy in large-scale protein database searches, contributes not only to scaling Blast but to scaling efficient search methods like MMSeqs2 and Diamond.This method supports any search tool that outputs results in the m8 file format, making it adaptable to new methods that also utilize this format. Incremental-Protein-Search offers the advantage of providing faster and more comprehensive retrieval by leveraging previous results, especially as the size of large-scale databases continues to grow.

## ACKNOWLEDGMENTS

This work is supported in part by funds from the National Science Foundation (#1936791, #1919691 and #2107108)

