## Supplementary materials for "iSeqsSearch: Incremental Protein Search for iBlast/iMMSeqs2/iDiamond"

### 1 Experimental Settings

1. Experiment Objective: To discern the differences between using an incremental method and not, by verifying that similar results can be achieved with the incremental approach.
2. Dataset Selection: The SCOPe Astral dataset[1] will be employed, comprising 305,543 sequences categorized into 7 SCOPe protein classes such as class a,b,c,d,e,f, and g. (astral-scopedom-seqres-gd-all-2.08-2023-01-06: latest version)

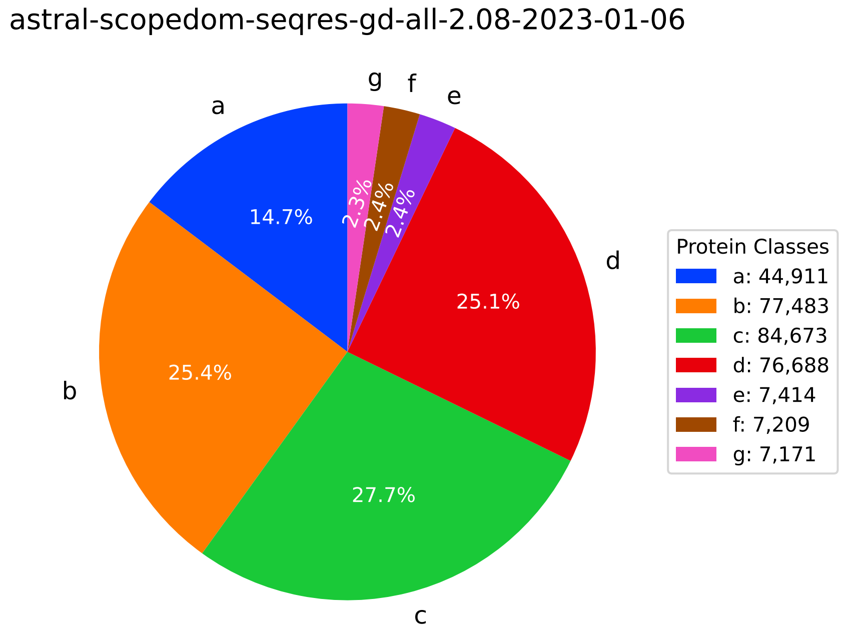


**Figure 1.** Protein Class Distribution

3. Experimental Design: The dataset will be divided into 10 batches. 1st batch is stratified random sampled and other 9 batches are random sampled based on protein class. A stratified randomly chosen 1st batch will serve as the query, while the remaining 9 batches will be sequentially combined for the search experiments. The experiments will be conducted using the default settings for BLASTP, MMseqs2, and DIAMOND. Only the number of threads was set to 32; all other settings were default.

### 2 SCOPe dataset

This project utilizes the SCOPe Astral dataset, which includes the extended database for the structural classification of proteins (SCOPe, Structural Classification of Proteins — extended). The features of this dataset are as follows: It consists of a total of 305,543 sequences from 4,327 species, categorized into 7 SCOPe protein classes, 1,257 protein folds, 2,065 protein superfamilies, and 5,084 protein families. The data, accessible via the SCOPe website, notably through the file 'astral-scopedom-seqres-gd-all-2.08-2023-01-06.fa', offers comprehensive information on protein structures and classifications, representing a rich resource for researchers exploring the vast diversity and complexity of biological systems.

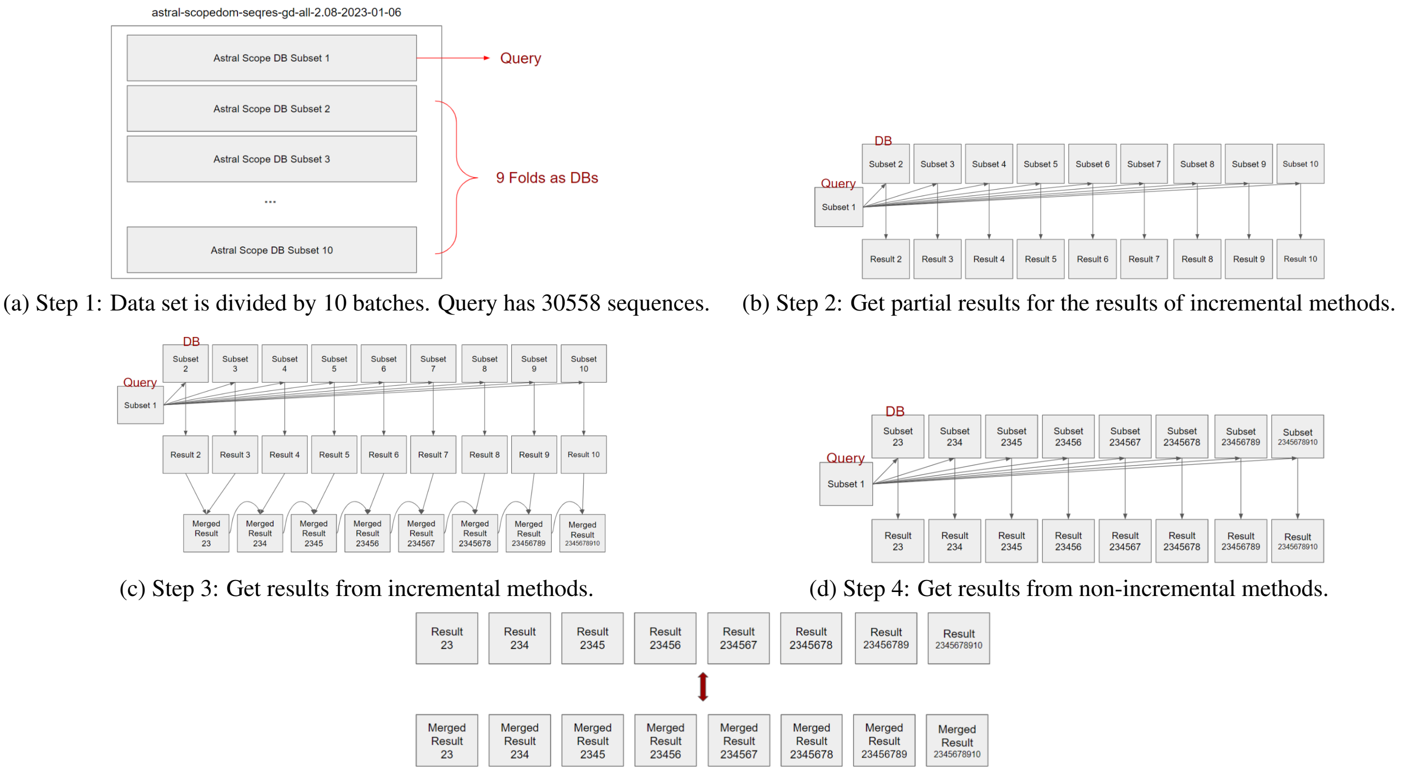


(a) Step 1: Data set is divided by 10 batches. Query has 30558 sequences.

(b) Step 2: Get partial results for the results of incremental methods.

(c) Step 3: Get results from incremental methods.

(d) Step 4: Get results from non-incremental methods.

(e) Step 5: Comparing incremental methods and non-incremental methods.

**Figure 2.** Illustration of the process for experiments

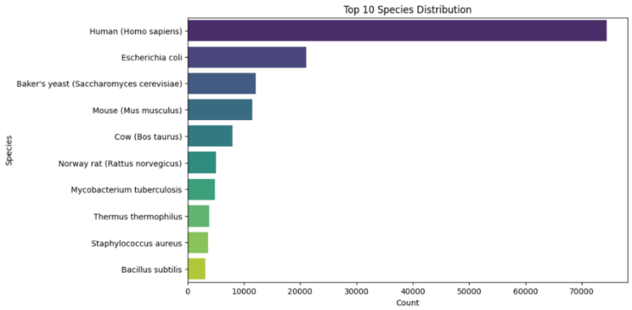


**Figure 3.** Species Distribution

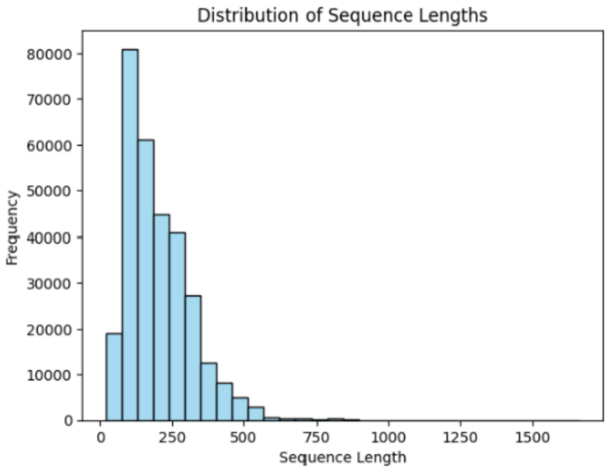


**Figure 4.** Sequence Length Distribution

### 3 Time Measure

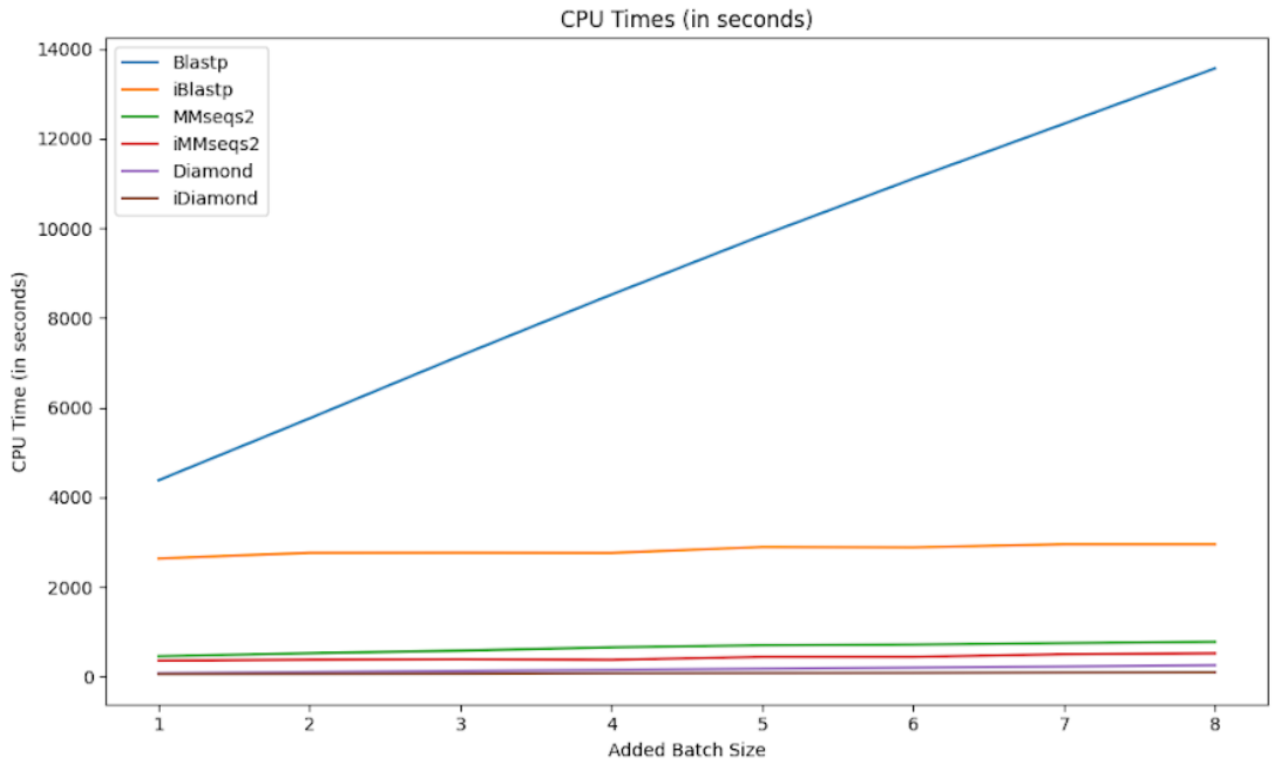


**Figure 5.** Time Comparision

The times for incremental methods represent the search time for the newly added batch and the time to merge these results with the previous results. For non-incremental methods, the times represent the search time for the entire database. The database consists of nine out of the ten portions of the entire Astral Scope dataset. In the experiments, one of the ten portions is used as the query, and the remaining batches from 2 to 10 are incrementally combined to test the search results.

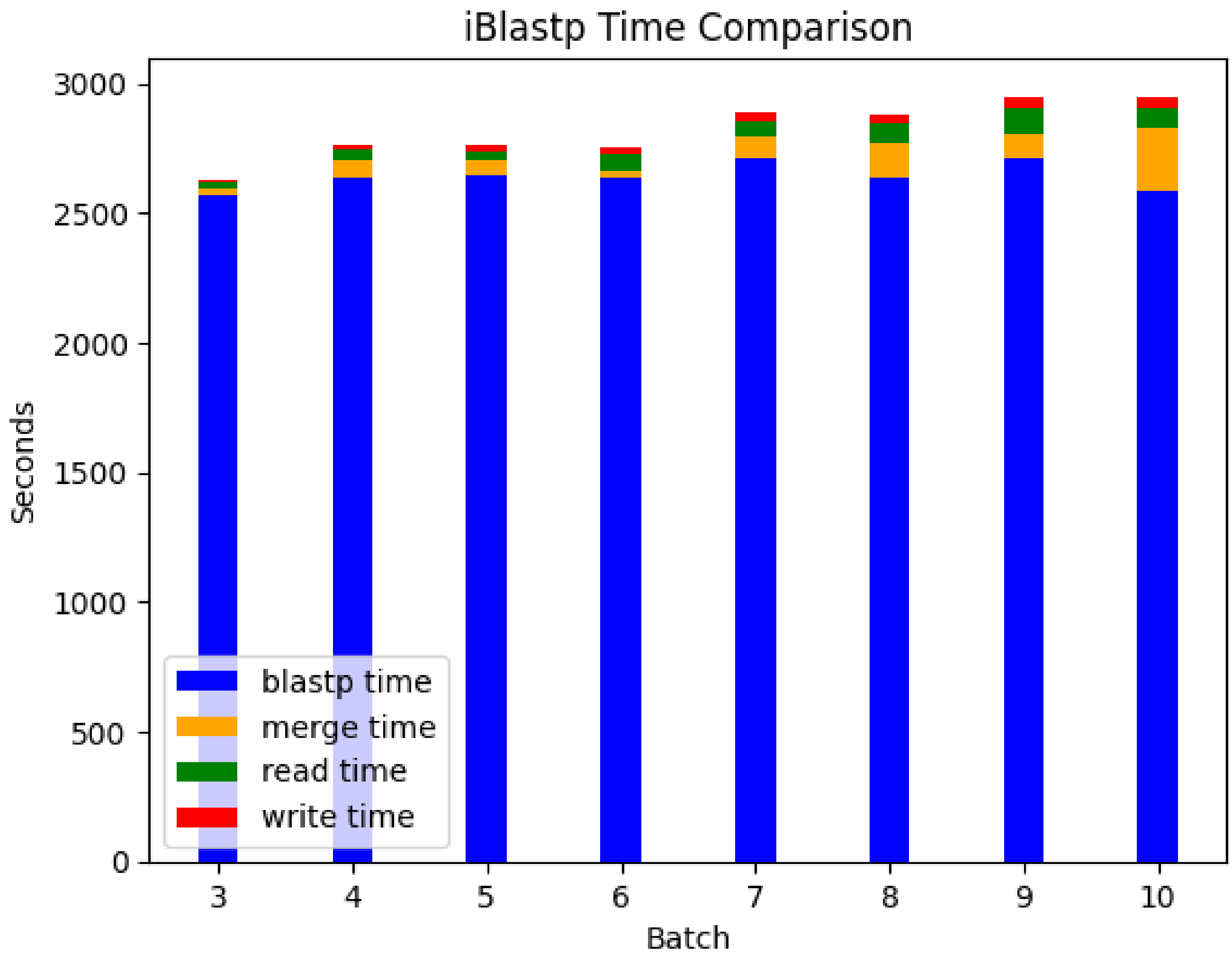


**Figure 6.** iBlast Time Comparison

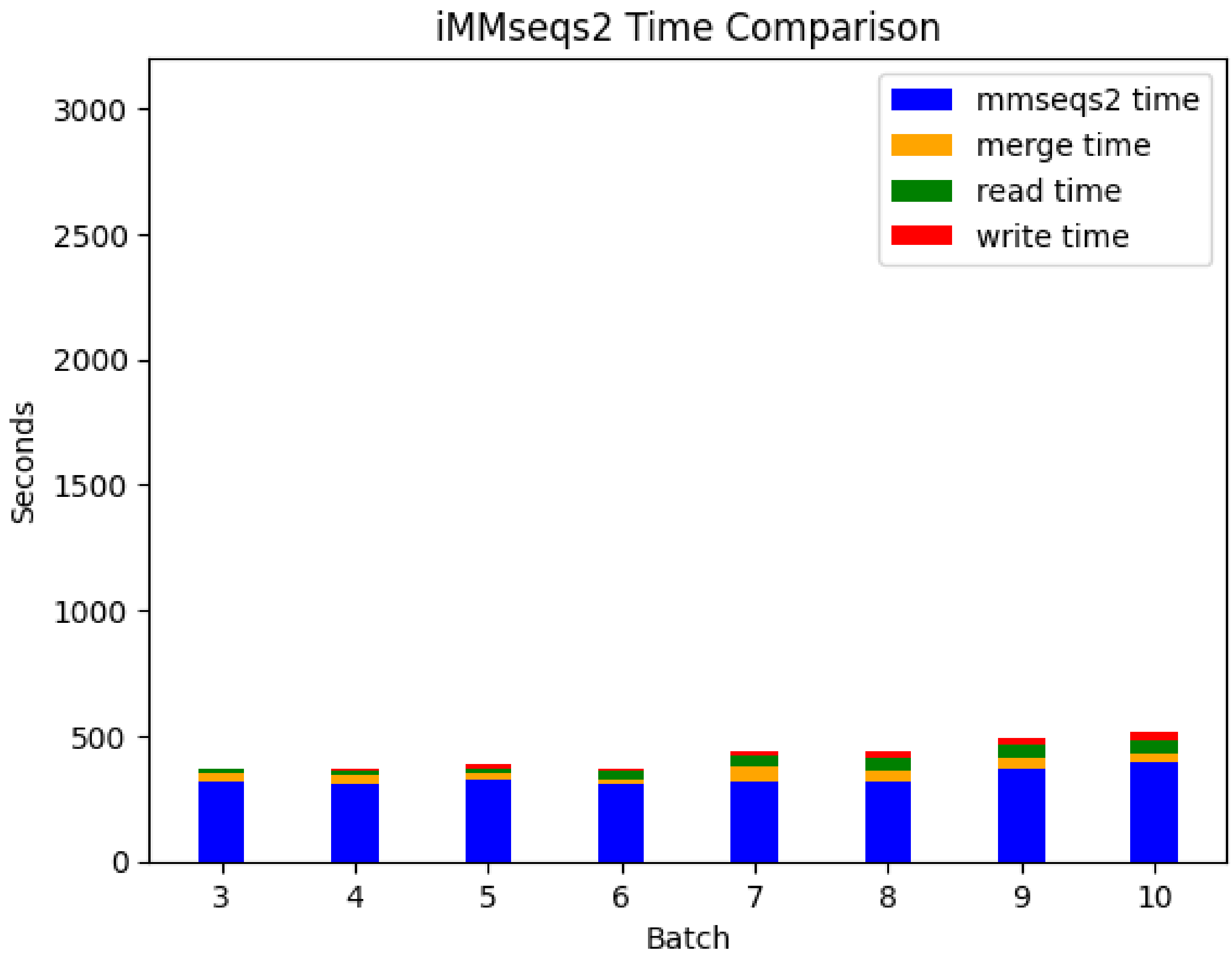


**Figure 7.** iMMseqs2 Time Comparison

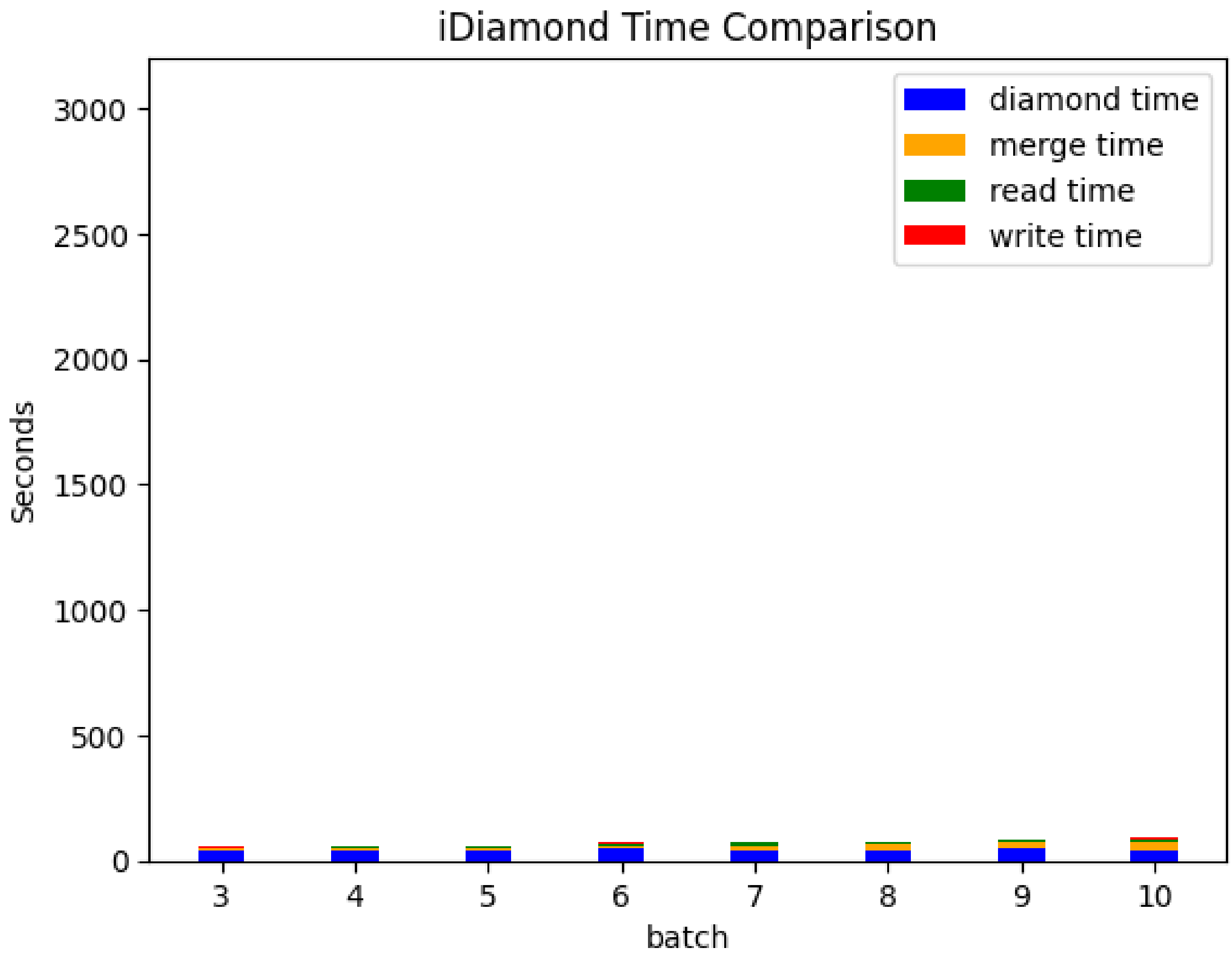


**Figure 8.** iDiamond Time Comparison

### 4 Protein SCOPe-Class Classification

We aimed to see how well we could classify the queries into their 7 classes (see Fig. 1) after the training of each random sub-batches. The graphs below compare the protein class f1 score of non-incremental methods and incremental methods based on the e-value top hit criterion. The results of the incremental experiments show a trend of increasing f1 score across all cases. The classification linearly increases as data is added in all cases up to 99% when all SCOPe[1] classes are known. In the last batch, every training class was included, leading to an increase in performance and achieving good results. The ratio of queries to the last batches is similar.

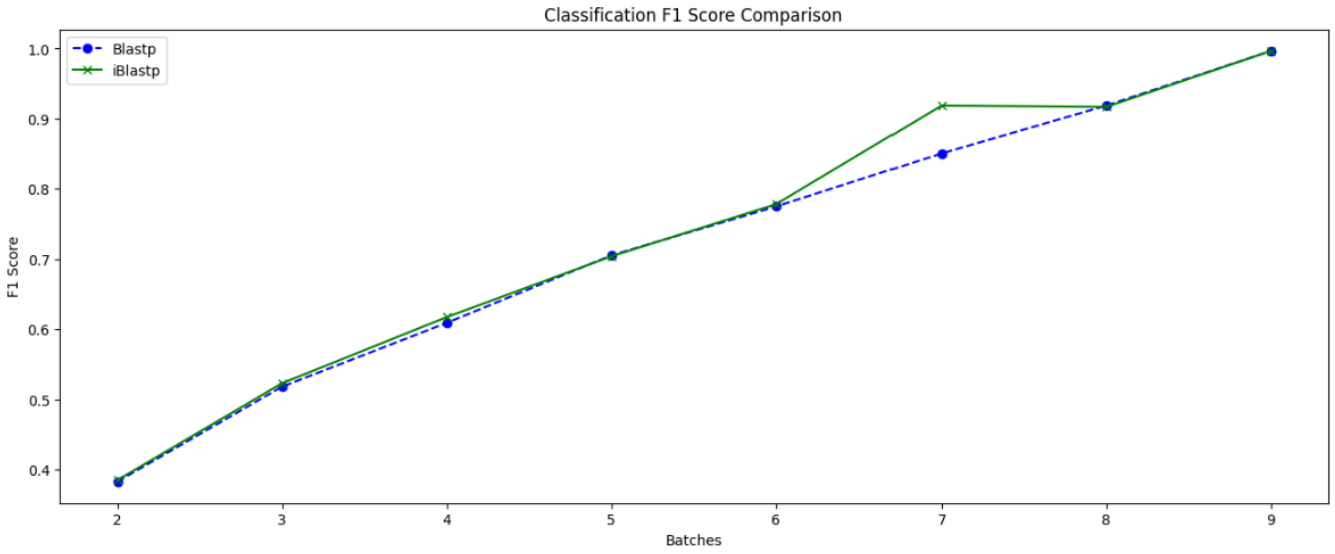


**Figure 9.** Blastp vs iBlastp

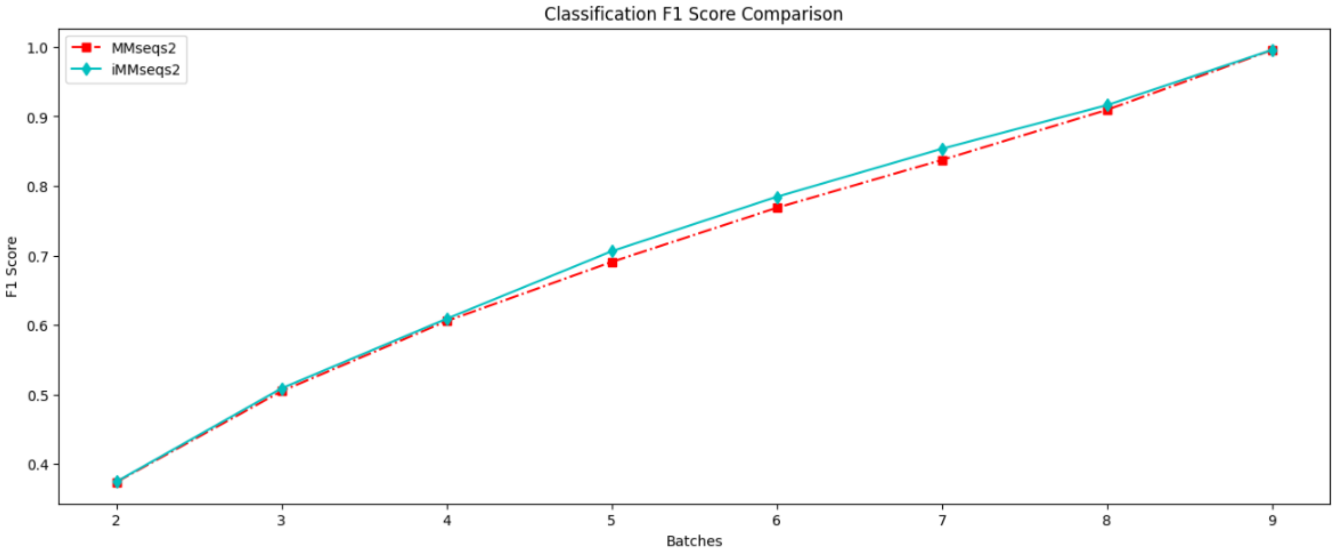


**Figure 10.** MMseqs2 vs iMMseqs2

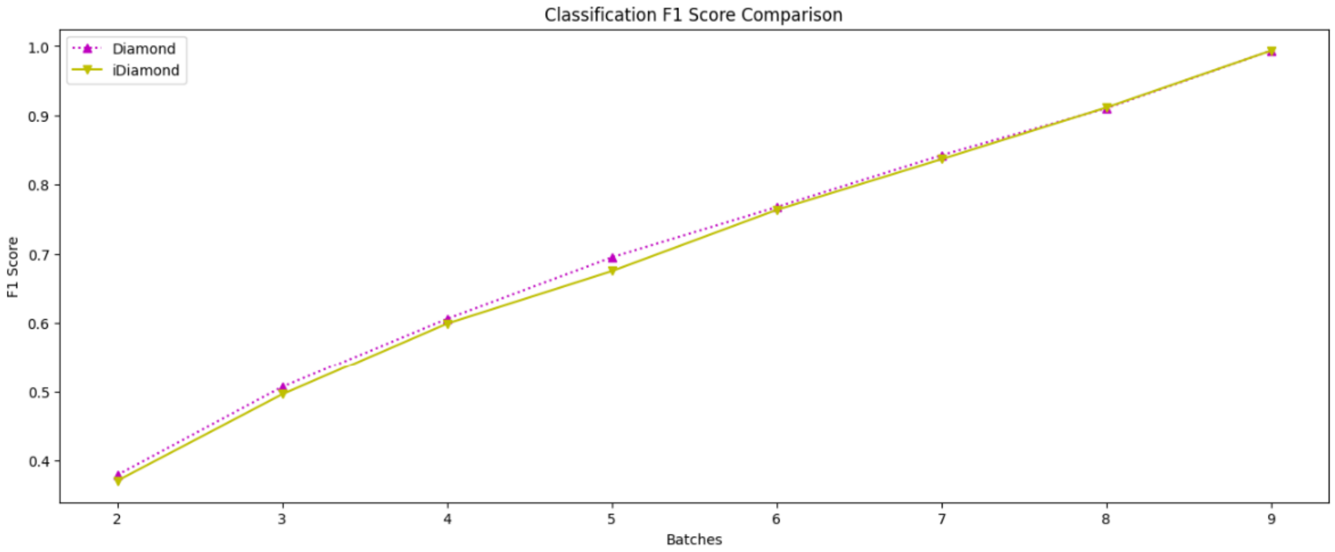


**Figure 11.** Diamond vs iDiamond

### 5 Venn Diagram with the hit number ratio

The Venn diagrams from the second and final batches illustrates that most of the hits from the non-incremental methods are included in the incremental methods. Specifically, the hits from Blastp are mostly included in the hits from iBlastp, and the hits from Diamond are also included in the hits from iDiamond. Similarly, the hits from MMseqs2 are included in the hits from iMMseqs2.

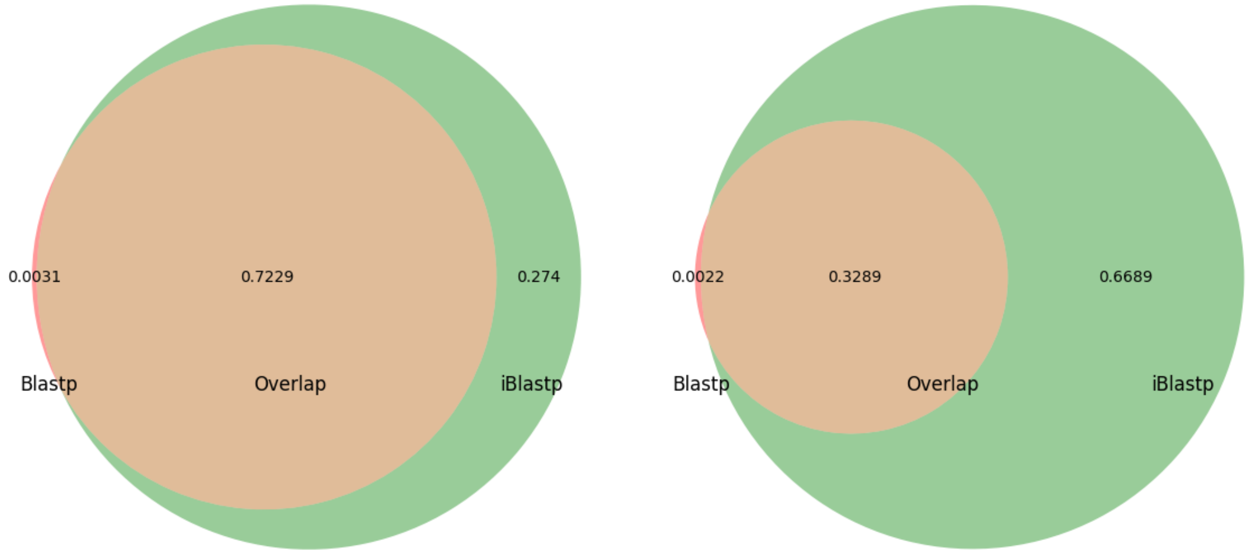


**Figure 12.** Blastp vs iBlastp with 2nd batch and last batch

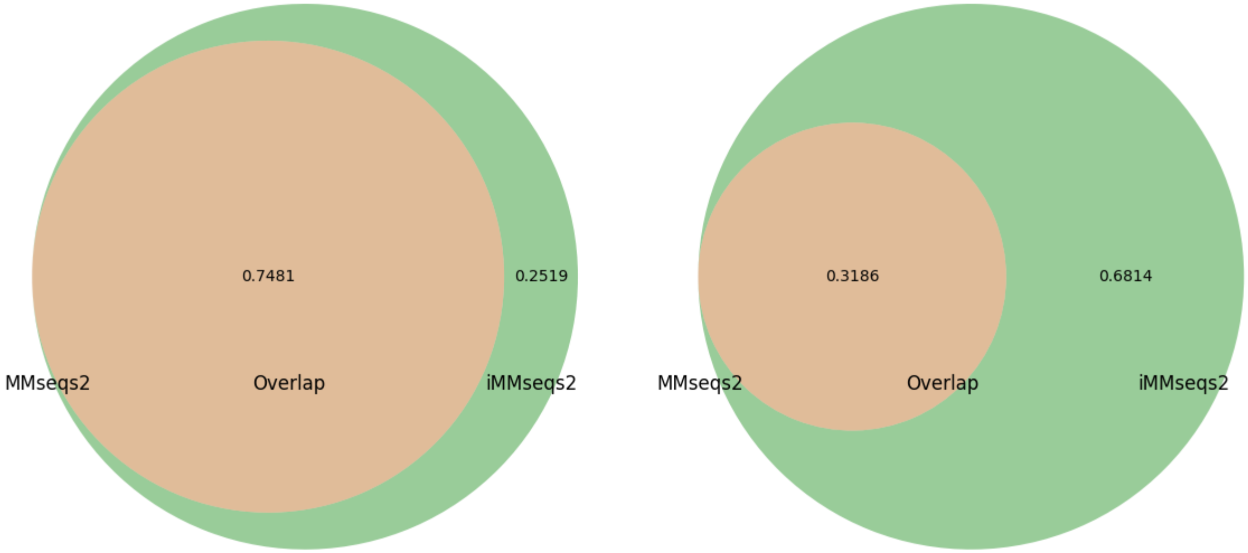


**Figure 13.** MMseqs2 vs iMMseqs2 with 2nd batch and last batch

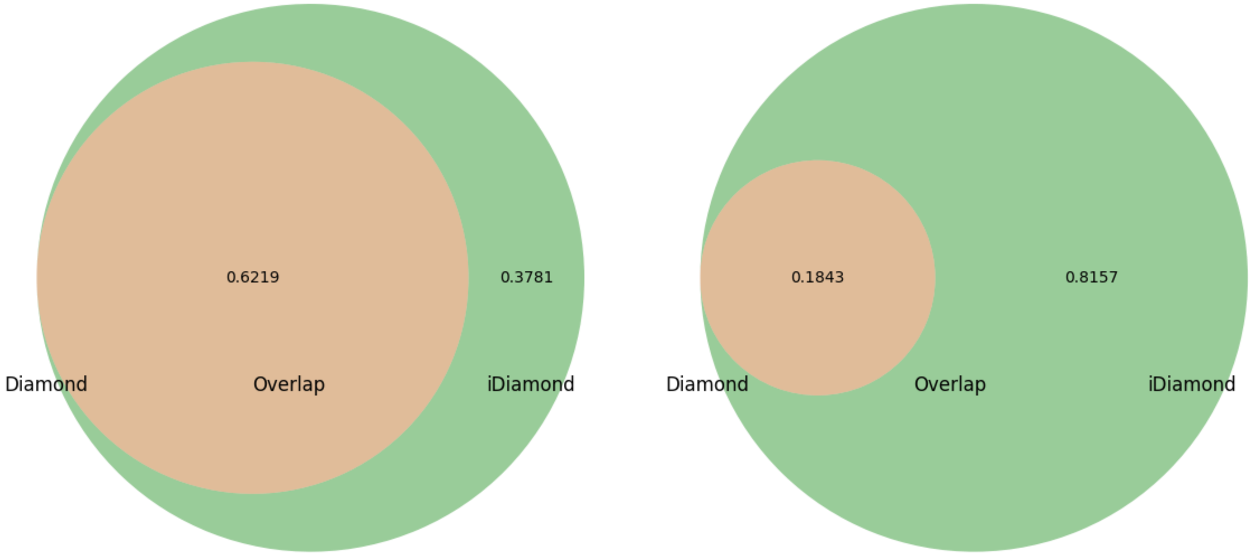


**Figure 14.** Diamond vs iDiamond with 2nd batch and last batch

### 6 No-hit per query ratio (non-stratified)

The figure below shows the proportion of queries that do not have a single hit as the search progresses incrementally. It can be seen that the incremental methods have a lower proportion of queries with no hits compared to the non-incremental methods. This indicates that incremental methods not only provide more hits but also cover a larger number of queries with hits than non-incremental methods. The first row at the top indicates the size of the increased batch.

| | 2 | 3 | 4 | 5 | 6 | 7 | 8 | 9 |
|---|---|---|---|---|---|---|---|---|
| Blastp | 0.0019 | 0.0018 | 0.0018 | 0.0013 | 0.0012 | 0.0012 | 0.0010 | 0.0010 |
| iBlastp | 0.0006 | 0.0001 | 0.0000 | 0.0000 | 0.0000 | 0.0000 | 0.0000 | 0.0000 |
| MMseqs2 | 0.0248 | 0.0170 | 0.0127 | 0.0095 | 0.0079 | 0.0067 | 0.0056 | 0.0049 |
| iMMseqs2 | 0.0247 | 0.0169 | 0.0124 | 0.0092 | 0.0076 | 0.0063 | 0.0053 | 0.0047 |
| Diamond | 0.0387 | 0.0266 | 0.0199 | 0.0154 | 0.0126 | 0.0106 | 0.0088 | 0.0075 |
| iDiamond | 0.0387 | 0.0265 | 0.0198 | 0.0154 | 0.0126 | 0.0106 | 0.0088 | 0.0075 |

**Figure 15.** This graph shows the ratio of no hit in query when batch increase

### 7 DCG measure

This table compares the DCG of incremental and non-incremental methods. The incremental method demonstrates superior DCG across all batches. The first row at the top indicates the size of the increased batch.

| | 2 | 3 | 4 | 5 | 6 | 7 | 8 | 9 |
|---|---|---|---|---|---|---|---|---|
| Blastp | 1275.4489 | 1611.0705 | 1886.7882 | 2117.4993 | 2327.6501 | 2513.8179 | 2677.9734 | 2829.0995 |
| iBlastp | 1381.0740 | 1854.8394 | 2296.3123 | 2716.6527 | 3122.8251 | 3516.9319 | 3901.4425 | 4277.6957 |
| MMseqs2 | 1304.7009 | 1618.7234 | 1876.5754 | 2088.6984 | 2270.1146 | 2417.2935 | 2544.9096 | 2658.6221 |
| iMMseqs2 | 1413.4290 | 1880.2749 | 2314.3664 | 2725.1164 | 3121.0765 | 3504.8746 | 3879.8322 | 4243.5533 |
| Diamond | 709.1630 | 774.2682 | 816.7293 | 847.9597 | 872.7081 | 891.9916 | 908.1241 | 922.0935 |
| iDiamond | 907.8142 | 1179.0332 | 1426.2112 | 1658.1434 | 1879.5196 | 2092.3212 | 2299.8423 | 2500.8473 |

**Figure 16.** Log DCG comparison for each batches

### 8 Case study(Protein family) : non-incremental method vs incremental method

The blast analysis has revealed duplicated hits. To ensure a fair comparison, particularly in the context of Blastp analysis, it is crucial to address these duplications and apply a stringent filter for E-values. In Blastp methodology, it has not removed these duplications nor filtered out hits with higher E-values. Therefore, to enhance the accuracy and reliability of the comparative analysis, we removed duplicated entries, retained only the hits with the smallest E-values, and discarded those with E-values greater than 1e-5, without limiting the number of hits. This adjustment will help maintain the integrity of our findings and ensure that the results are both robust and meaningful.

Query case ’d2ap2c1(right label: b.1.1.1)’ and ’d6iyia_(a.1.1.0)’ are used for this case study. One protein was randomly selected from the SCOPe alpha protein class and another from the beta protein class. These proteins were chosen to ensure diverse structural representation and to assess the algorithm’s performance across different protein folds.

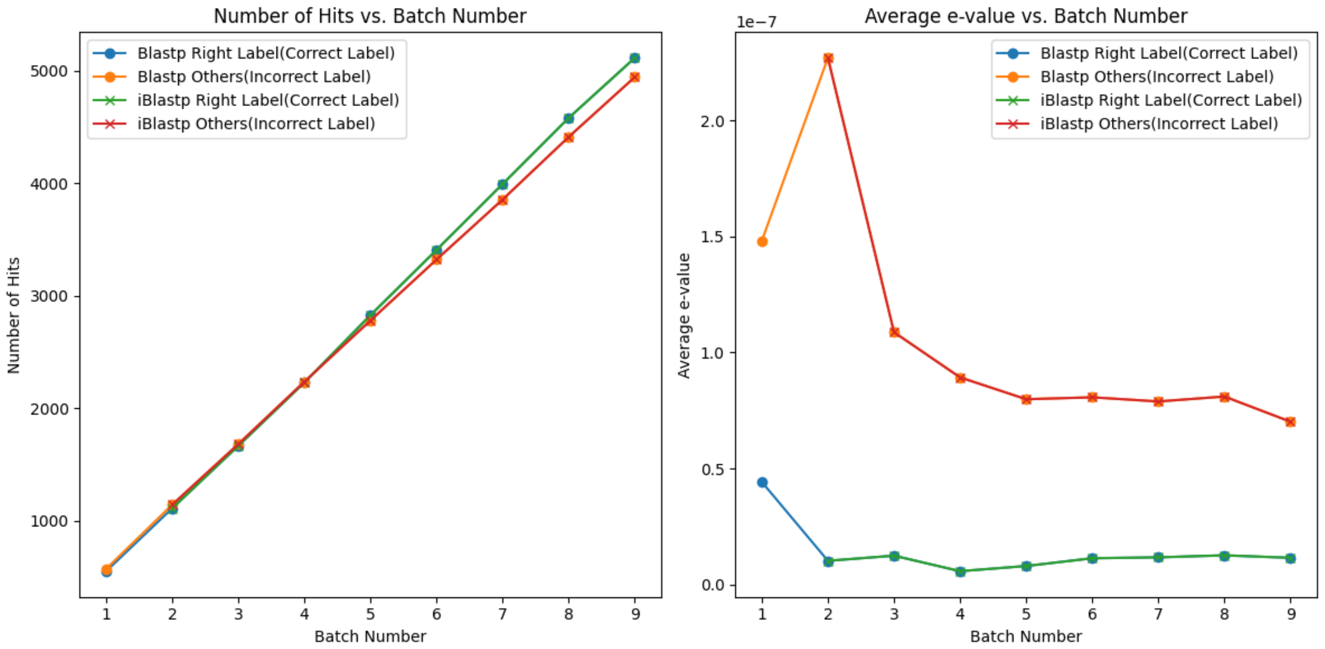


**Figure 17.** Query "d2ap2c1" Case Study: Blastp vs iBlastp where “Right label" is the Correct Protein Family label and others mean all incorrect labels.

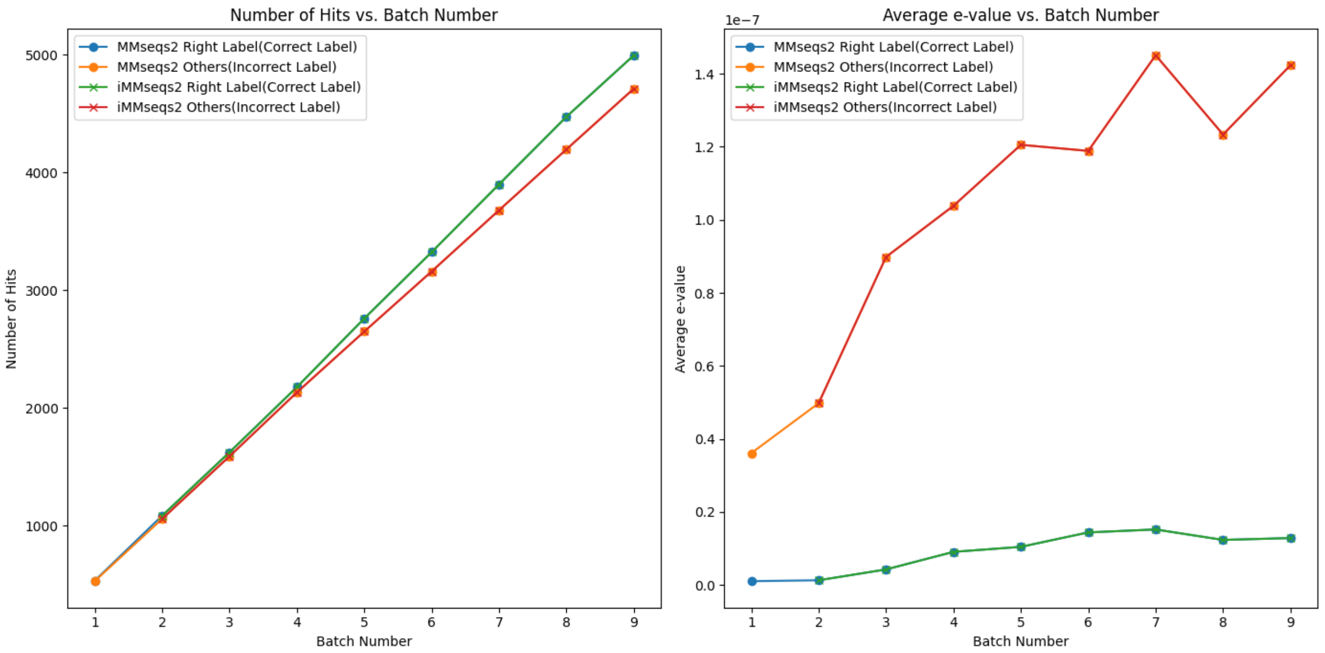


**Figure 18.** Query "d2ap2c1" Case Study: MMseqs2 vs iMMseqs2 where “Right label" is the Correct Protein Family label and others mean all incorrect labels.

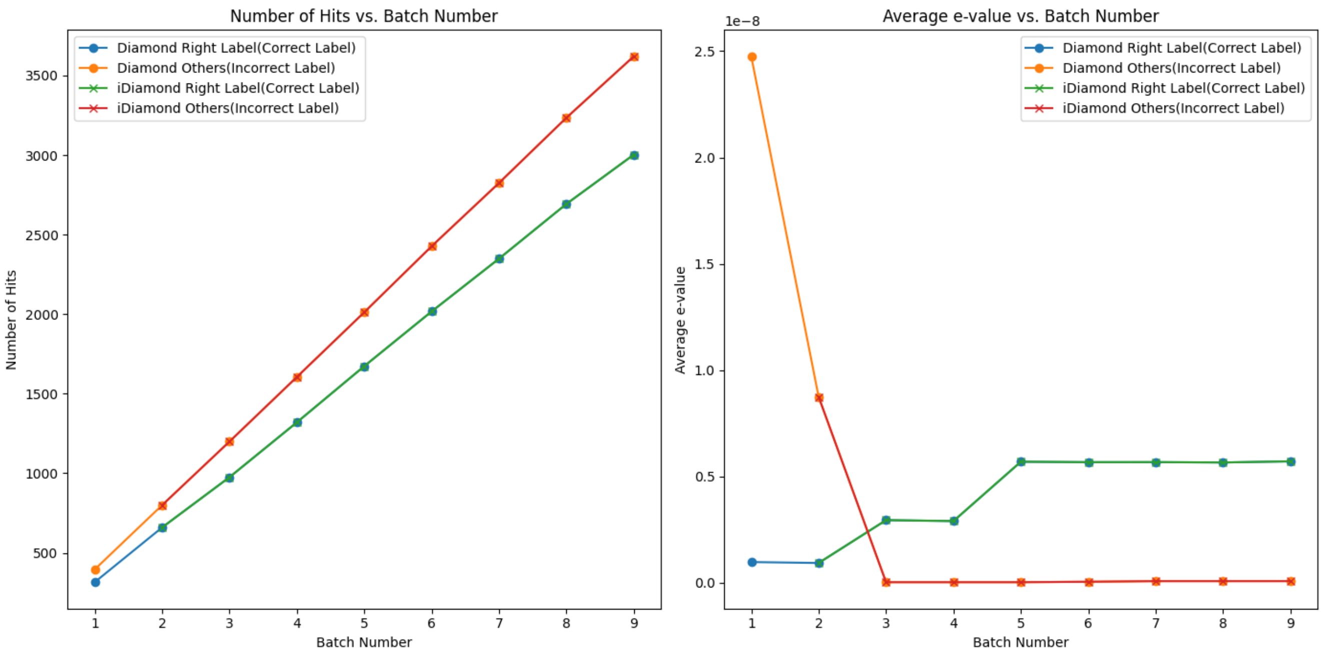


**Figure 19.** Query "d2ap2c1" Case Study: Diamond vs iDiamond where “Right label" is the Correct Protein Family label and others mean all incorrect labels.

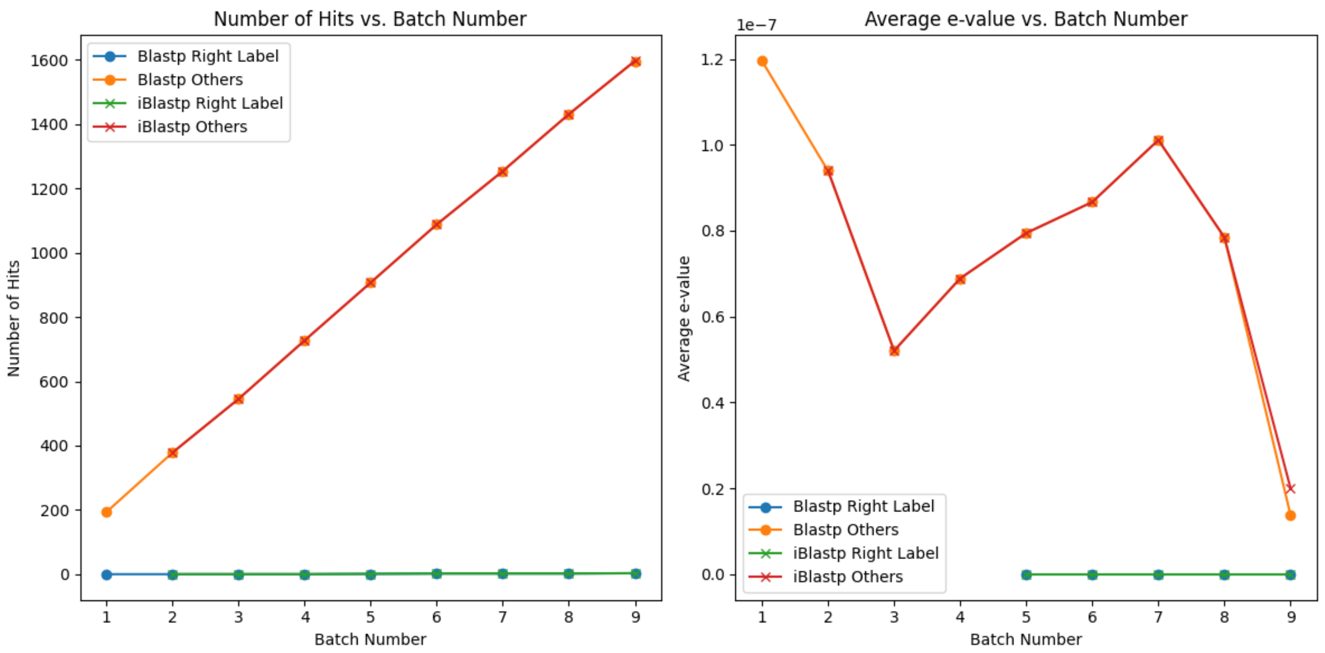


**Figure 20.** Query "d6iyia_" Case Study: Blastp vs iBlastp where “Right label" is the Correct Protein Family label and others mean all incorrect labels.

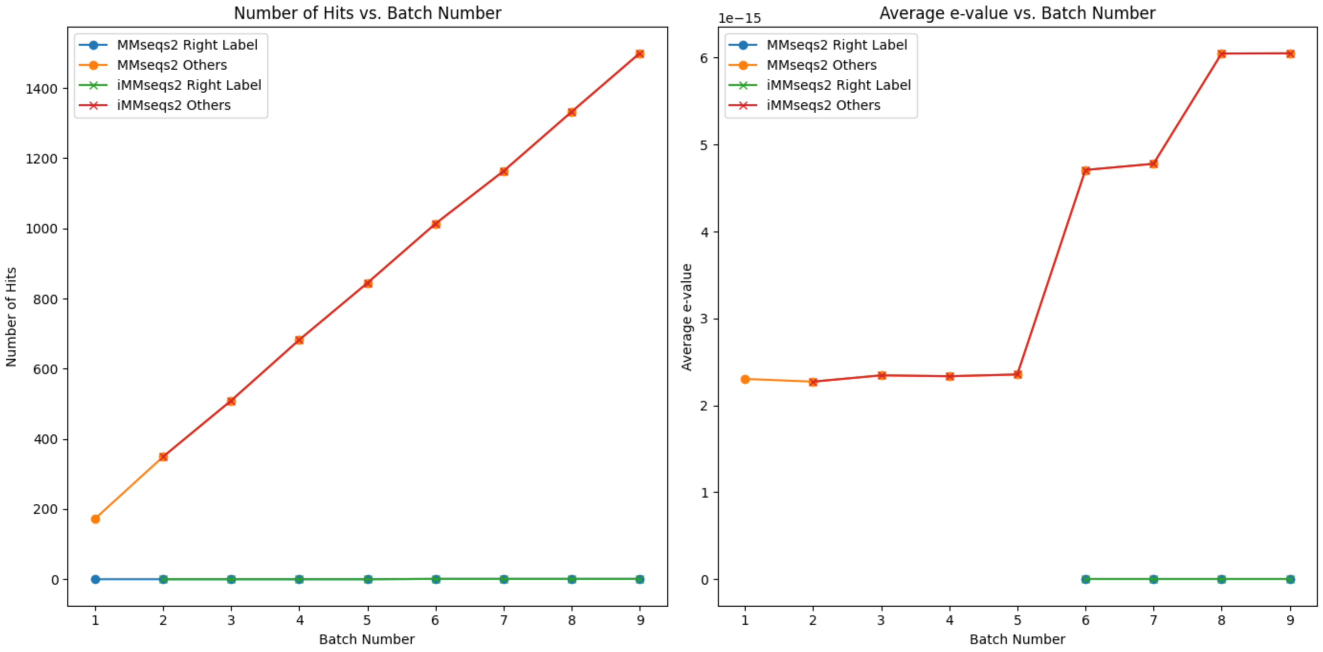


**Figure 21.** Query "d6iyia_" Case Study: MMseqs2 vs iMMseqs2 where “Right label" is the Correct Protein Family label and others mean all incorrect labels.

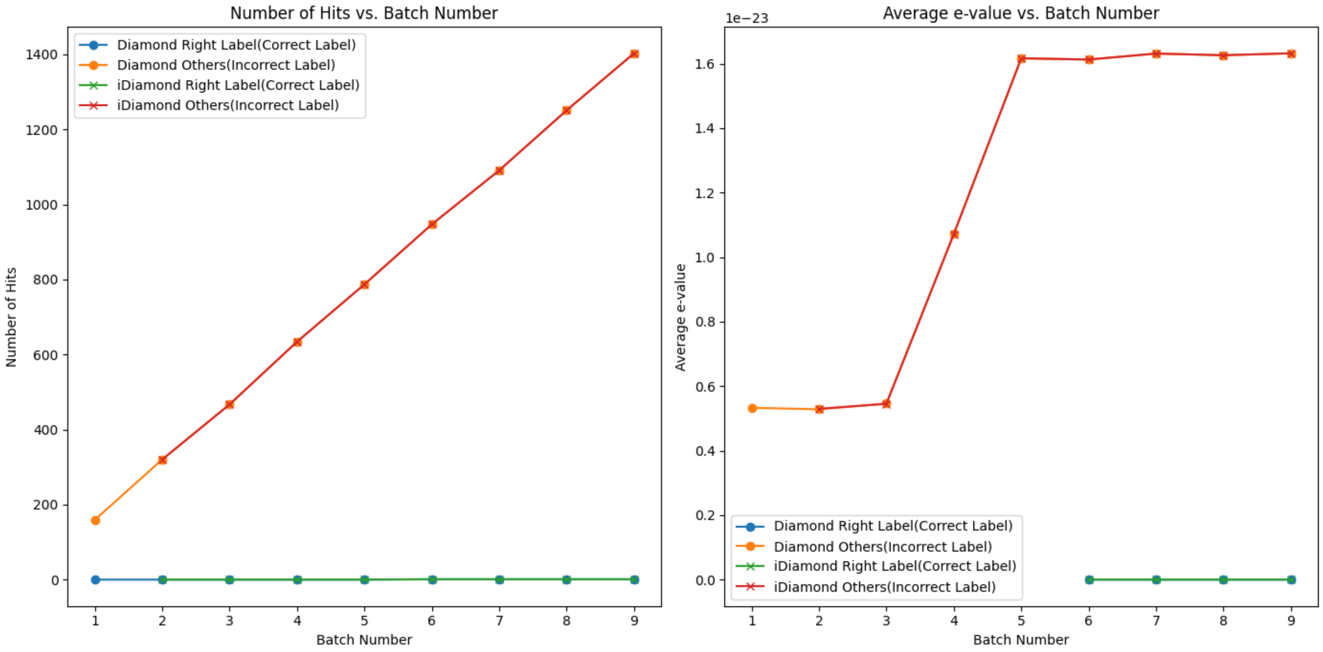


**Figure 22.** Query "d6iyia_" Case Study: Diamond vs iDiamond where “Right label" is the Correct Protein Family label and others mean all incorrect labels.

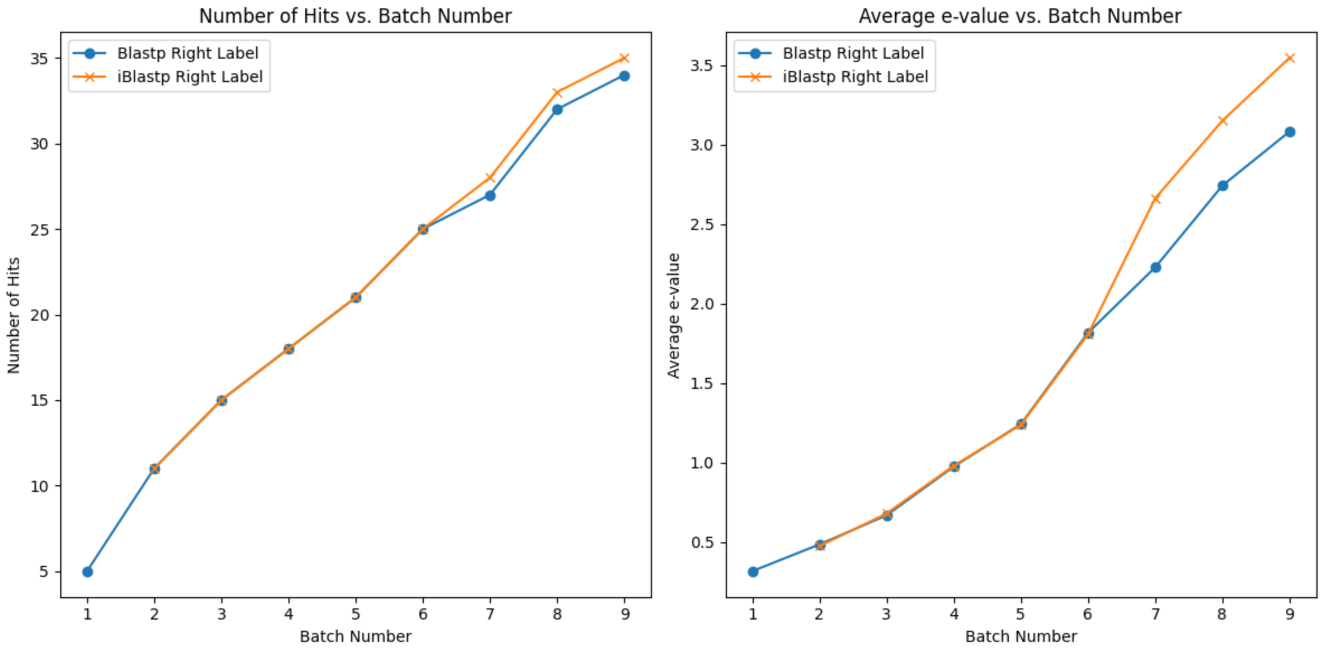


**Figure 23.** Query "d6iyia_" Case Study: Blastp vs iBlastp without 1e-5 threshold where “Right label" is the Correct Protein Family label.

This experiment is conducted with specific conditions. The hit number limit was removed, and duplicates were handled by retaining the hit with the smallest e-value, excluding results with an e-value greater than or equal to 1e-5. The resulting graph above reflects these conditions. The number of hits between incremental methods (iBlastp, iMMseqs2 and iDiamond) and non-incremental methods (Blastp, MMseqs2, Diamond) remained similar even as the batch size increased. The average e-value also showed very similar results. Additionally, without 1e-5 threshold, iBlastp shows more hits and higher average e-value compared to Blastp like Figure 23.
